# Botswana’s wildlife losing ground as Kalahari Wildlife Management Areas (WMAs) are dezoned for livestock expansion

**DOI:** 10.1101/576496

**Authors:** Derek Keeping, Njoxlau Kashe, Horekwe (Karoha) Langwane, Panana Sebati, Nicholas Molese, Marie-Charlotte Gielen, Amo Keitsile-Barungwi, Quashe (/Uase) Xhukwe, !Nate (Shortie) Brahman

**Author notes:** DEREK KEEPING (Corresponding author) Department of Renewable Resources, University of Alberta, 751 General Services Building, Edmonton, Alberta, T6G 2H1, Canada (and) Latisa Tracking Solutions, Maun, Botswana. orchid.org/0000-0001-5050-2226. deceased (!Nate (Shortie) Brahman).

## Abstract

Botswana is rightfully lauded for maintaining 42% of its land base for conservation, and ranking top in the world for effort to conserve megafauna. Yet during the recent preservationist political climate characterized by a hunting moratorium and deterioration of CBNRM, changes to the national land use map suggest Botswana is losing grip on its lofty status. While conservation attention is turned to elephants and KAZA, discussion about the official dezoning of 8,268 km^2^ of free-ranging wildlife estate in the Kalahari ecosystem has been notably absent. Using track-based methods we quantified wildlife populations inhabiting this relinquished wilderness now awaiting imminent conversion to fenced and private livestock holdings. We find that the affected areas contain approximately 3,900 free-ranging large herbivores and 50 large carnivores, all of which will become consumed, displaced or potential conflict animals. Erstwhile publicly-owned wildlife particularly important for local communities will effectively become de facto private property for an elite minority. The land use changes spell negative consequences for wildlife not only via mechanisms of habitat loss and edge effects but also reduced landscape connectivity between protected areas that limits seasonal movements and gene flow thus eroding long-term population resilience in a drought-prone environment. As Botswana’s agricultural lobby continues to exert pressure on the Kalahari ecosystem, we suggest that ground surveys conducted by Kalahari trackers be implemented to inform decision-making rather than relying on the inadequate coarse-grained aerial survey record only.

---

> To those devoid of imagination a blank place on the map is a useless waste; to others, the most valuable part. (Leopold, 1949, p. 176)
>
> The livestock industry operates within a kind of conceptual Dark Age…in ignorance of the ecological matrix from which their meat mountains and milk quotas derive. Their activities expand at inordinate cost to the long-term health of African environments and natural resources. (Kingdon, 1997, p. 323)

Botswana’s Kgalagadi Transfrontier Park (KTP) and Central Kalahari Game Reserve (CKGR), together with the rangelands connecting these two great Kalahari protected areas, comprise what is possibly the largest wildest ecosystem remaining south of the Sahara (WCS and CIESIN, 2005) (Fig. 1). In significant contrast to the northern and eastern parts of Botswana criss-crossed with veterinary fences which cordon off the Kalahari and severely disrupt wildlife movements beyond (Williamson & Williamson, 1984), no such barriers to wildlife movement exist within the free-ranging Kalahari ecosystem outlined in Figure 1. Conceivably, a large-bodied animal can move between Two Rivers in the southwest and Kuke corner in the northeast - a straight-line distance approaching 700 kilometres - without encountering hindrance to its travel. This distance is much greater than that over which famed migrations in East Africa, and that recently noted as the “longest in Africa” (Naidoo et al., 2016) occur.

**FIG. 1.**
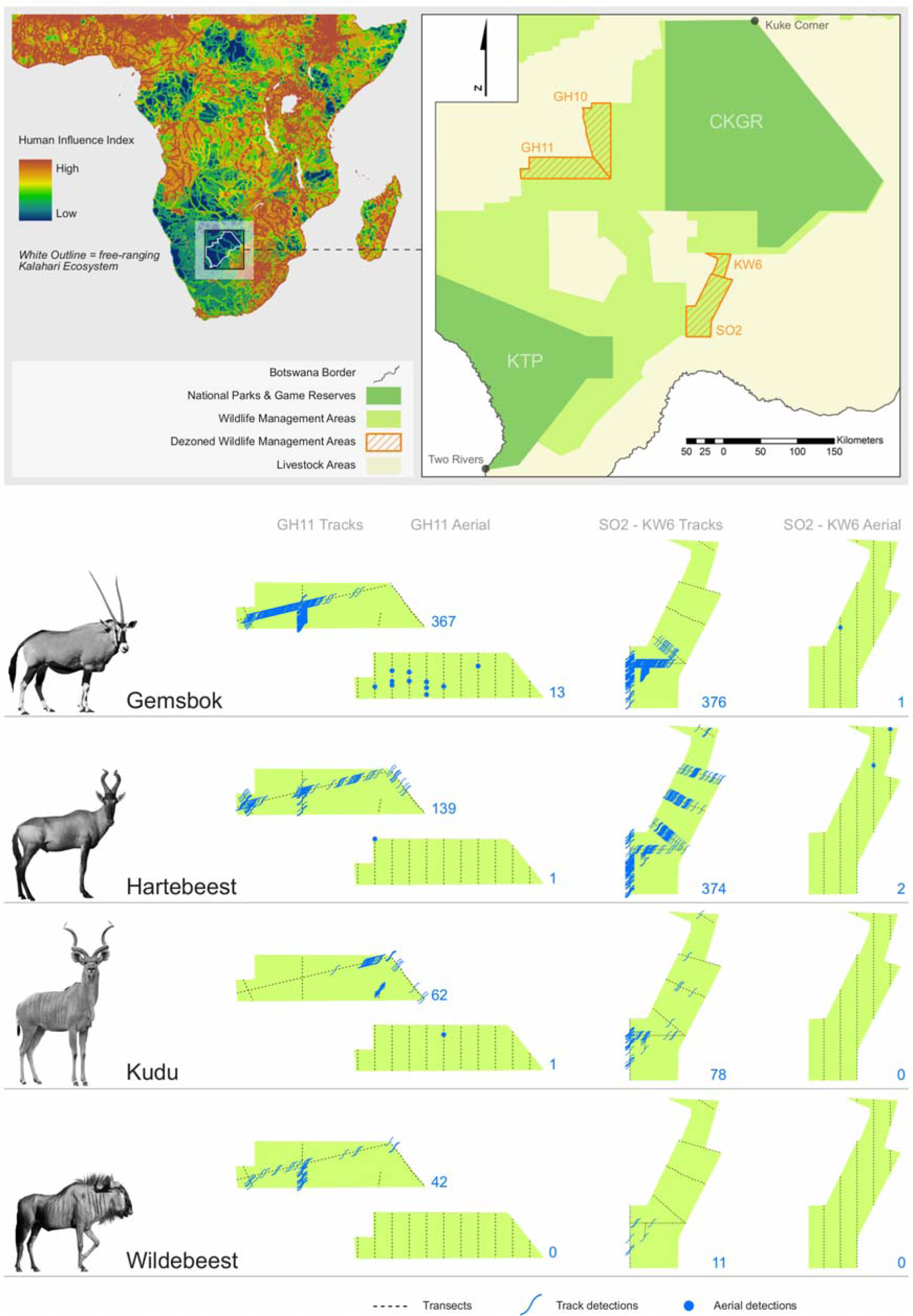
Locations and numbers of track VS aerial survey detections of large antelope species in three dezoned Wildlife Management Areas in relation to the larger free-ranging Kalahari ecosystem and human influence (dataset from WCS and CIESIN, 2005) in sub-Saharan Africa. Aerial detections are combined 2012 and 2015 surveys, while track detections are likewise from twice repeated sampling of GH11 transects, but SO2-KW6 differ in that track transects were sampled once only.

Not the two parks but the space in between them, referred to as the Schwelle, has long been recognized as the core area for semi-migratory wildlife (DHV, 1980; Bonifica, 1992). Yet foundational reports hardly few decades old stressing the importance of these areas to the long-term viability of free-ranging wildlife populations seem to have been quickly forgotten, and Verlinden’s (1997) speculative “populations of [free-ranging antelopes] in the southern and central Kalahari might become separated when the area occupied by livestock increases…between the two areas” uncritically accepted to have taken place already. It is now a well entrenched belief in Botswana that wildlife has lost access to the Schwelle and become largely restricted to protected areas (e.g. Selebatso et al., 2018).

We suggest Botswana’s aerial survey record plays prominent in this jump to conclusions; both its inherent limitations, and its interpretation. In the absence of other data, there is an over-reliance on aerial surveys to inform spatial density-distributions of large herbivores. But coverage is poor: 96-97% of the Kalahari landscape is never surveyed. Coarsely-resolved map outputs therefore display many blank pixels deceptively unoccupied by wildlife (Keeping et al., 2018). The second source of influential data available is telemetry collaring. During the early 1990’s, VHF-collared wildebeest moved long distances between CKGR and the Schwelle (Bonifica, 1992). Since then, limited GPS collaring efforts, including the most recent (Selebatso et al., 2018), have failed to validate trans-Kalahari movements. As a research approach GPS telemetry is practically limited to infinitesimal sample sizes and thus weak population-level inferences (Hebblewhite & Haydon, 2010), but perhaps the usual precautionary interpretation demanded in such instances is relaxed when scant other data exist.

Tragically, this recent underrating of Kalahari WMAs is now threatening to become a self-fulfilling prophecy. Botswana’s 2014 hunting moratorium coincided with the first substantial change in WMA boundaries since their inception in 1986. Four long-gazetted WMAs (GH10, GH11, SO2, KW6) comprising 8,268 km^2^ were officially dezoned to allow privatized fenced livestock expansion (Fig. 1). Although approved on paper, these land use changes have yet to be implemented on the ground. We ground-truthed GH11, SO2 and KW6 at opportunistic periods between 2014-17, conducting track surveys along convenient linear features (infrequently driven sand trails and cutlines) that bisect the affected areas. Two expert local observers seated over the front of a 4×4 vehicle enumerated all large wild herbivore and carnivore track interceptions that endured weathering enough to be identifiable, estimated track age to the nearest 24 hr period, and recorded each interception with GPS.

We applied the FMP formula (Stephens et al., 2006) to estimate average wildlife densities from track counts within the dezoned areas. By “average” we imply the estimates are derived from data collected over 2014-17, as opposed to a snapshot in time. The FMP formula describes true animal density by linking track interceptions (≤ 24 hrs) over a known transect distance to average day range of the population creating the tracks. Empirical day range estimates were available for large carnivores, gemsbok and wildebeest from previous studies (see Keeping 2014). For species lacking empirical day range data (hartebeest, ostrich, kudu), we used allometrically-derived day ranges and for all species followed the bootstrapping procedures described in Keeping (2014) to generate density estimates with confidence intervals.

GH10 was not surveyed but we assumed similar wildlife densities in the other dezoned areas and extrapolated population estimates for the total dezoned area. This assumption is fair considering GH10 has similar habitat and edge to GH11, but also potentially conservative given the closer proximity of GH10 to CKGR and lack of linear features for vehicle access. For comparison with track data, we extracted the most recent aerial survey records from 2012 and 2015 and estimated populations in the dezoned areas using Jolly’s method II (ratio method) for unequal-sized sample units (Jolly, 1969). For comparisons of detections between tracks and aerial survey, we utilized the complete track data set (i.e. tracks that were fresh but also multiple days old), while track-based density calculations utilized only a subset of those data (i.e. tracks ≤ 24 hrs old).

Contrasting the paucity of aerial survey detections, tracks disclosed a more complete picture of wildlife inhabiting the dezoned WMAs (Fig. 1). We estimated 3,878 large herbivores, the most numerous being disturbance-sensitive gemsbok, and 53 large carnivores, most ranking Vulnerable or worse on the IUCN Red List, occupying the affected areas and thus directly impacted by changing land use (Table 1). Our estimates fail to account for edge effects that will extend beyond the dezoned areas after land use changes are implemented. Wildlife densities are already depressed within the dezoned areas due to edge effects from adjacent livestock areas, and these dezoned WMAs presently function as buffers to higher wildlife densities in the core WMAs. In the Kalahari, fenced ranches are not hard boundaries; they tend to be grossly overstocked and owners release cattle into adjacent areas when grazing is depleted inside (Darkoh & Mbaiwa, 2001). Thus, once the land use changes are implemented, the area of affected wildlife habitat and numbers of animals impacted will be potentially much greater than presented in Table 1. Beyond habitat loss and edge effects, the configuration of landscape loss is concerning. The southern and most direct connection (SO2-KW6) between KTP and CKGR will be lost entirely, leaving a single narrowed linkage remaining between the two parks (Fig. 1). The GH11 and GH10 dezonings sever a wildebeest migration route along fossil drainages utilized during below average rainfall years (Bonifica, 1992), undermining recovery of the depressed Kalahari population.

**TABLE 1.**
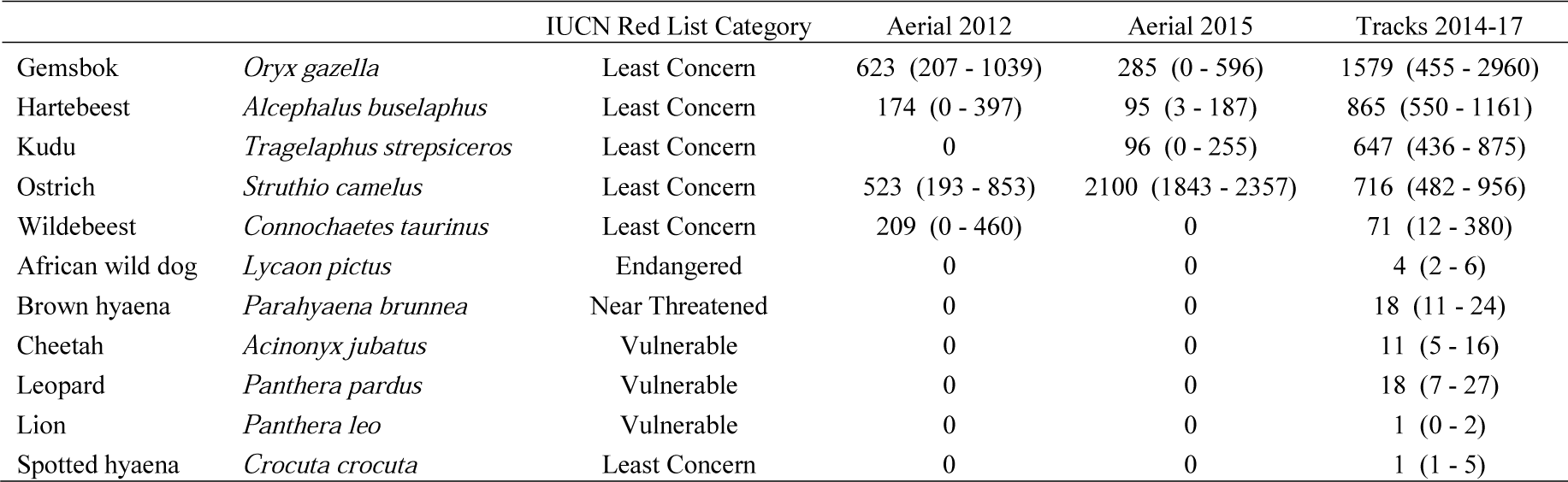
Population estimates (95% confidence limits) from two aerial surveys and an averaged track survey of large herbivore and carnivore species inhabiting Kalahari Wildlife Management Areas (8,268 km^2^) dezoned for fenced livestock expansion

Whilst Botswana’s aerial survey reveals valuable insights at the countrywide or regional scale, at the scale of dezoned WMAs it returned too few detections to be very useful. The sampling rate in the Kalahari is too low (3-4% area coverage), and secondly thresholds of detection from the air may exist at low wildlife densities. Population estimates varied markedly between years due to the stochastic nature of limited sightings (Table 1). Contrasting the incomplete snapshot provided by direct sightings, the time-integrated nature of track counts ensured high detection rates (i.e. detecting animals that had moved over greater distances than aerial strip widths). Such data are more informative in two respects: high detection rates improve distribution maps, and larger sample sizes encourage more accurate density estimation. We therefore urge collaboration with expert local trackers to assess wildlife populations in vulnerable habitats in Botswana before decisions to relinquish such habitats for livestock expansion are made.

In the Kalahari environment, cattle expansion and intensification degrades rangelands (Dougill et al., 2016), negatively impacts carbon balance (Thomas, 2012), exacerbates inequality and extreme poverty (Chanda et al., 2003), and precludes tourism development by mutual exclusion of wildlife and wildlife-based livelihoods. Botswana’s cattle economy makes up less than 2% total GDP (Statistics Botswana, 2018), while tourism, predominantly wildlife and wilderness based, contributes approximately 11.5% (WTTC, 2018). Despite this, the loss of wildlife estate continues to be framed as an inevitable “compromise” to the cattle industry (Department of Lands, 2009). The boundaries of Botswana’s WMAs can change at any time as provided by the Wildlife Conservation and National Parks Act (DWNP, 1992). As a result, formally gazetted WMAs comprising a substantial area of wildlife habitat have been dezoned wholescale, without an Environmental Impact Assessment process. The ease of this event does not bode promising for the future of free-ranging wildlife in this largest, as yet unrealized wilderness asset in southern Africa.

## Author contributions

Study design and fieldwork: DK, NK, HL, PS, NM, QX and !NB; data analysis and writing the article: DK, MG and AK.

## Acknowledgements

We are grateful to the Government of Botswana via Department of Wildlife and National Parks and Ministry of Environment, Natural Resources Conservation and Tourism for permissions to conduct research. We thank Comanis Foundation for supporting our research and Antje Hellwig for help with graphics. MG thanks F.R.S-FNRS Fonds de la Recherche Scientifique for funding.

## Ethical standards

All authors have abided by the Code of Conduct for contributors. The research involved non-invasive wildlife surveys, therefore ethics and animal welfare approvals were not required.

## References

Bonifica (1992) Technical assistance to the project: initial measures for the conservation of the Kalahari ecosystem. Final Report under EDF project No. 6100.026.14.001 to the Department of Wildlife and National Parks, Government of Botswana. Bonifica, Rome, Italy.

Chanda, R., Totolo, O., Moleele, N., Setshogo, M. & Mosweu, S. 2003. Prospects for subsistence livelihood and environmental sustainability along the Kalahari Transect: The case of Matsheng in Botswana’s Kalahari rangelands. Journal of Arid Environments, 54(2), 425–445.

Darkoh, M.B.K. & Mbaiwa, J.E. (2001) Sustainable development and resource conflicts in Botswana. In African pastoralism: Conflicts, institutions and government (eds M.A.M. Salih, D. Dietz & A.G.M. Ahmed), pp. 39–55. Pluto Press, London, UK.

Department of Lands (2009) Review of National Land Use Map. Final Report. Ministry of Lands and Housing, Government of Botswana, Gaborone, Botswana.

Department of Wildlife and National Parks (1992) Wildlife Conservation and National Parks Act. Chapter 38:01. Government of Botswana, Gaborone, Botswana.

DHV (1980) Countrywide Animal and Range Assessment Project. Final Report to Ministry of Commerce and Industry & Department of Wildlife and National Parks, Government of Botswana. DHV Consulting Engineers, Amersfoort, Netherlands.

Dougill, A.J., Akanyang, L., Perkins, J.S., Eckardt, F.D., Stringer, L.C., Favretto, N., et al. (2016) Land use, rangeland degradation and ecological changes in the southern Kalahari, Botswana. African Journal of Ecology, 54(i1), 59–67.

Hebblewhite, M. & Haydon, D.T. (2010) Distinguishing technology from biology: a critical review of the use of GPS telemetry data in ecology. Philosophical Transactions of the Royal Society B: Biological Sciences, 365(1550), 2303–2312.

Jolly, G. M. (1969) The treatment of errors in aerial counts of wildlife populations. East African Agriculture and Forestry Journal: special edition, 34, 50–55.

Keeping, D. (2014) Rapid assessment of wildlife abundance: estimating animal density with track counts using body mass–day range scaling rules. Animal Conservation, 17(5), 486–497.

Keeping, D., Burger, J.H., Keitsile, A.O., Gielen, M.C., Mudongo, E., Wallgren, M., et al. (2018) Can trackers count free-ranging wildlife as effectively and efficiently as conventional aerial survey and distance sampling? Implications for citizen science in the Kalahari, Botswana. Biological Conservation, 223, 156–169.

Kingdon, J. (1997) The Kingdon Field Guide to African Mammals. Academic Press, London, UK.

Leopold, A. (1949) A Sand County Almanac: And Sketches Here and There. Oxford University Press, New York, USA.

Naidoo, R., Chase, M.J., Beytell, P., Du Preez, P., Landen, K., Stuart-Hill, G., et al. (2016) A newly discovered wildlife migration in Namibia and Botswana is the longest in Africa. Oryx, 50(1), 138–146.

Selebatso, M., Bennitt, E., Maude, G. & Fynn, R.W. (2018) Water provision alters wildebeest adaptive habitat selection and resilience in the Central Kalahari. African Journal of Ecology, 56(2), 225–234.

Statistics Botswana (2018) Gross Domestic Product: Second Quarter of 2018. http://www.statsbots.org.bw [accessed 03 January 2019].

Stephens, P.A., Zaumyslova, O.Y. Miquelle, D.G., Myslenkov, A.I. & Hayward, G.D. (2006). Estimating population density from indirect sign: track counts and the Formozov-Malyshev-Pereleshin formula. Animal Conservation, 9(3), 339–348.

Thomas, A.D. (2012) Impact of grazing intensity on seasonal variations in soil organic carbon and soil CO2 efflux in two semiarid grasslands in southern Botswana. Philosophical Transactions of the Royal Society of London B: Biological Sciences, 367(1606), 3076–3086.

Verlinden, A. (1997) Human settlements and wildlife distribution in the southern Kalahari of Botswana. Biological Conservation, 82(2), 129–136.

Wildlife Conservation Society, and Center for International Earth Science Information Network - Columbia University. (2005) Last of the Wild Project, Version 2, 2005 (LWP-2): Last of the Wild Dataset (Geographic). http://dx.doi.org/10.7927/H4348H83. [accessed 15 September 2018].

Williamson, D. & Williamson, J. (1984) Botswana’s fences and the depletion of Kalahari wildlife. Oryx, 18(4), 218–222.

World Travel & Tourism Council (2018) Travel & Tourism Economic Impact 2018 Botswana. https://www.wttc.org [accessed 03 January 2019].

